# Analysis of multistage in vitro fertilization data with mixed multilevel outcomes using joint modelling approaches

**DOI:** 10.1101/173534

**Authors:** Jack Wilkinson, Andy Vail, Stephen A Roberts

**Affiliations:** Centre for Biostatistics, Institute of Population Health, Manchester Academic Health Science Centre (MAHSC), University of Manchester, Manchester, UK; Research and Development, Salford Royal NHS Foundation Trust, Salford M6 8HD, UK; Centre for Biostatistics, Institute of Population Health, Manchester Academic Health Science Centre (MAHSC), University of Manchester, Manchester M13 9PL, UK

**Keywords:** in vitro fertilisation, joint modelling, mixed data, multilevel modelling, multistage treatment data, multivariate responses

## Abstract

In vitro fertilization comprises a sequence of interventions concerned with the creation and culture of embryos which are then transferred to the patient’s uterus. While the clinically important endpoint is birth, the responses to each stage of treatment contain additional information about the reasons for success or failure. Joint analysis of the sequential responses is complicated by mixed outcome types defined at two levels (patient and embryo). We develop three methods for multistage analysis based on joining submodels for the different responses using latent variables and entering outcome variables as covariates for downstream responses. An application to routinely collected data is presented, and the strengths and limitations of each method are discussed.

## 1. Background and motivation

In vitro fertilization (IVF) is a complex multistage procedure for the treatment of subfertility. Typically, a ‘cycle’ of IVF begins with the administration of drugs to stimulate the patient’s ovaries and promote the release of oocytes (eggs). The oocytes are collected from the patient and are then fertilised either by mixing or injecting them with sperm. The resulting embryos are cultured for several days. Finally, one or more of the best embryos are selected for transfer to the woman’s uterus, where it is hoped that they will implant and develop into a healthy baby. Treatment may fail at any stage of the cycle (if no oocytes are recovered from the ovaries, no good quality embryos are produced, or those transferred do not implant), in which case the subsequent stages are not undertaken.

The sequential nature of IVF means that the patient’s response can be measured at each stage of the treatment (Heijnen et al. 2004): the stimulation of the ovaries can be evaluated by the number of oocytes collected; the fertilization and culture stages can be evaluated by the number and quality of embryos produced; and the success of the transfer procedure can be evaluated according to whether or not a child is born as a result. Figure 1 displays a schematic of the IVF cycle. A recent review of outcome measures used in IVF RCTs showed that there is considerable interest in these ‘intermediate’ or ‘procedural’ outcomes of IVF; 361 distinct numerators were identified, and the median (IQR) number of distinct outcomes reported per trial was 11 (7 to 16) (Wilkinson et al. 2016).

**Figure 1:**
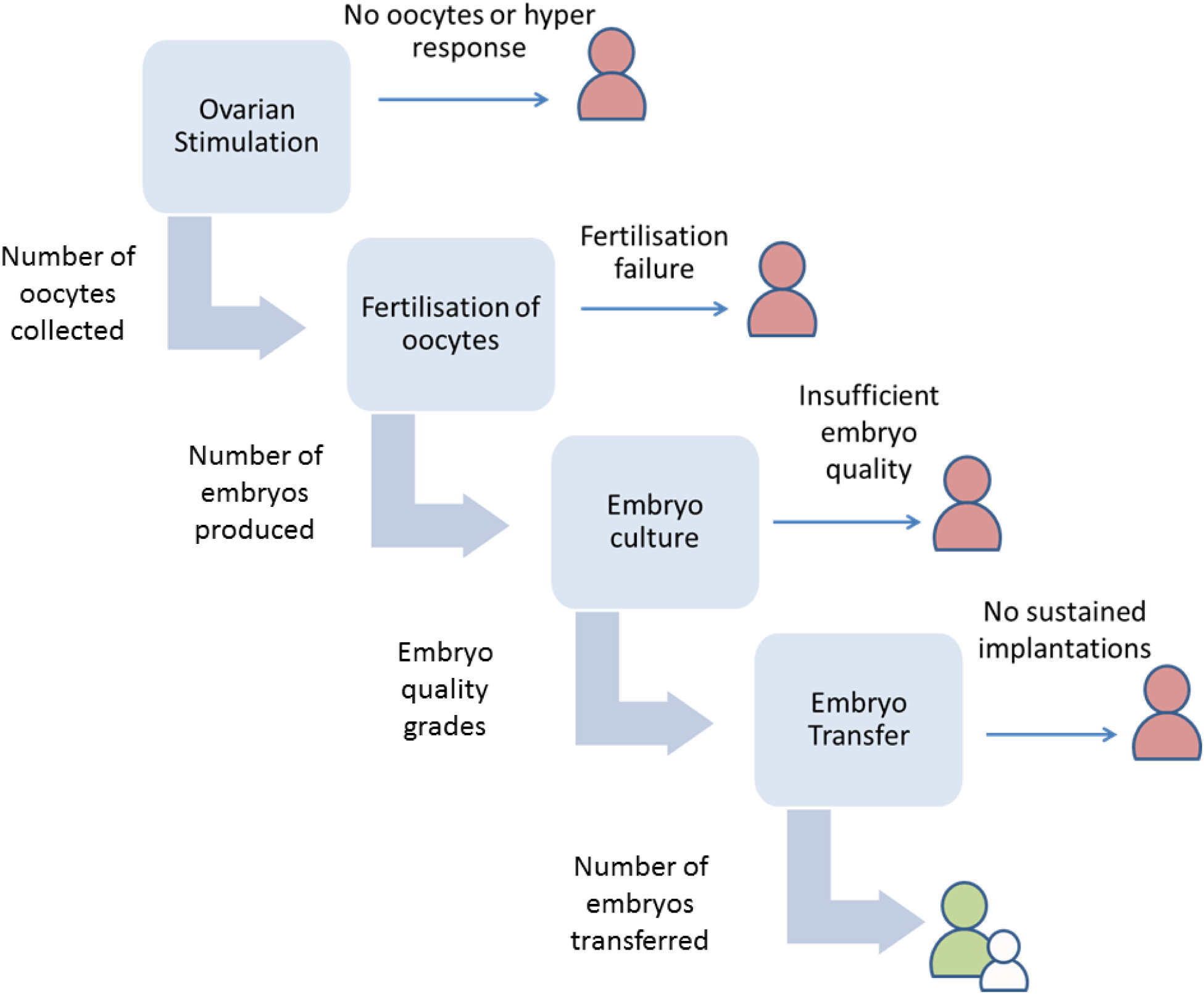
Schematic of the IVF cycle.

The interest in procedural outcomes in IVF research is not surprising. While the most relevant measure of success for patients is the birth of a child (Min et al. 2004, Heijnen et al. 2004, Legro et al. 2014), establishing the effects of treatments and patient characteristics on procedural outcomes might increase our mechanistic understanding of how IVF works and how it might be improved. The question of how outcomes at each stage of the process relate to one another also appears to be relevant to designing and evaluating IVF interventions. In response, two approaches for the analysis of multistage IVF data have recently been proposed (Maity et al. 2014, Penman et al. 2007). The first is a discrete time-to-event approach that treats the stages of the IVF cycle as a series of ‘failure opportunities’ (Maity et al. 2014). Each woman’s response data then comprise a vector of binary indicator variables denoting whether they failed at this stage, or proceeded to the next. The second treats the stage of the cycle reached by the patient as an ordinal response, and models this using continuation ratio regression (Penman et al. 2007). Both of these approaches allow us to answer research questions relating to the effects of baseline treatment and patient characteristics on IVF response, while preserving the sequential nature of the data. Both share similar limitations, however. In particular, both treat the responses at each stage as dichotomous ‘success or failure’ events. This wastes a great deal of information, since it is more informative to measure the number of oocytes obtained from the ovaries than merely whether a sufficient quantity were available to enable the cycle to continue; and it is more informative to measure the quality of any embryos obtained than merely whether there were any available for transfer. These methods are also incapable of accommodating outcomes defined at different levels of a multilevel structure; some outcomes (eg: number of oocytes) may be defined for each patient, while others (eg: embryo quality) are defined for the patient’s individual embryos. In addition, while these methods allow for differential effects of covariates at each stage through the inclusion of interaction terms, they do not allow for different covariates to be included as predictors for the different stage-specific responses.

While methods for the analysis of sequential IVF data exist therefore, it remains to identify techniques capable of incorporating the variety of outcome types encountered in this setting, and moreover responses which are defined at different levels of a two-level data structure (embryos and patients). This includes counts of oocytes, ordinal embryo quality scales, binary birth indicator variables, and so on. Methods for the analysis of multivariate responses of mixed outcome types are hardly new (eg: Goldstein 2003) but have received considerable attention in recent years (see de De Leon and Chough (2013) for a comprehensive collection of the state of the art). While much of this work has focussed on the joint analysis of time-to-event and longitudinal response data (see reviews by Gould et al. (2015) and Tsiatis and Davidian (2004)), approaches capable of accommodating different combinations of outcome types have been described (McCulloch 2008; Dunson 2000, Gueorguieva 2001, Gueorguieva and Agresti 2001, Dunson and Herring 2005, Dunson et al. 2003, Goldstein et al. 2009). Typically, these involve the inclusion of shared (McCulloch 2008, Dunson 2000, Gueorguieva 2001, Dunson 2000, Dunson and Herring 2005) or otherwise correlated (Gueorguieva and Agresti 2001, Goldstein et al. 2009) latent variables in ‘submodels’ for the different response variables. These latent variables accommodate dependency between the response variables in the model. Moreover, by estimating the parameters governing the distribution of these latent variables, we can examine both the direction and degree of association between a patient’s responses. A further attractive feature of latent variable approaches is that they can be used to jointly model responses measured at different levels of a multilevel data structure (Goldstein et al. 2009, Dunson et al. 2003). These methods do not appear to have been discussed in the context of multistage treatments however.

Given the strict temporal ordering of the stages, an alternative strategy for the analysis of IVF data would be to explicitly model the relationships between the patient’s stage-specific responses using a series of conditional regression equations (Blalock 1961). Under this sort of approach, each response variable would be included as a covariate in the regression equations for each of the subsequent, or ‘downstream’, responses. An advantage of these approaches is that they allow direct and indirect effects of the procedural responses on downstream outcomes to be distinguished (Pearl 2001). A third strategy we might consider would be to combine the two approaches hitherto described, and simultaneously link submodels for each response using latent variables while including the response variables as covariates in the downstream response models. This would then resemble the endogeneous treatment models employed in the econometrics literature (Terza 1998), or multiprocess models that have been employed in education research (Steele et al. 2009).

In this paper, we develop methodology for the analysis of multistage IVF data, with mixed response types (count, ordinal, and binary) defined at different levels of a two-level data structure (patients and embryos). We describe three approaches in which distinct submodels are used for the various response variables. In the first, we include correlated latent variables and estimate the relationships between the responses. In the second, we adopt an outcome regression approach where response variables enter into regression equations for downstream response variables as covariates. This approach can be considered as a set of separate regression models. In the third, we consider an endogenous response model where we combine both of these approaches, by including upstream response variables as covariates in downstream submodels, and also allowing the submodel-specific latent variables to be correlated. The remainder of the manuscript is structured as follows. In section 2, we describe the models. In section 3 we illustrate the use of the methods with an application to a routine clinical database. This is followed by a discussion in section 4. We conclude with some brief recommendations in section 5.

## 2. Models

Here we describe latent variable, outcome regression and endogenous response modelling approaches to the analysis of multistage IVF data. The approaches have several key features in common. First, they all include distinct submodels for each of the response variables considered in the cycle. We include six response variables for patient *j* = 1, … *n* and their embryos *i* = 1,…,n_j_, and hence six submodels, in the current presentation: the number of oocytes (eggs) obtained from ovarian stimulation (a county 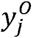); the fertilisation rate when the oocytes are mixed with sperm 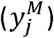 two measures of embryo quality (cell evenness and degree of fragmentation 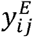 and 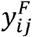 both measured using ordinal grading scales); an indicator denoting whether one or two embryos were transferred to the patient (denoted by a binary variable 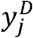 ) and another 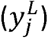 indicating whether or not the transfer of embryos resulted in the live birth of one or more babies (a live birth event, or LBE) (Fig.1). These are listed in temporal order, with the exception of the two embryo quality scales, which are coincident. We include the decision to transfer two embryos (known as double embryo transfer, or DET) in the model because it is an important predictor of transfer success which is partially determined by the outcomes of the earlier stages. A second feature common to the approaches is that once a patient has dropped out of the cycle, they do not appear in the submodels corresponding to the downstream responses. In the following, we ignore the possibility that each patient may undergo multiple cycles of IVF, noting that the models could be extended to three levels (embryos nested within cycles nested within women) by adding additional random scalar terms (Goldstein 2011).

### 2.1 Correlated latent variable approach

This approach requires the use of latent variable representations for the various submodels constituting the larger model. Each patient *j* has associated vectors of responses 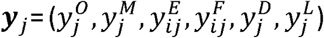 and Of underlying latent variables 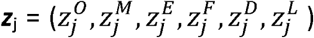. Both of these vectors may be partially observed due to drop-out or outright failure before completion of the treatment. We then posit a multivariate Normal distribution for the latent variables, and estimate the elements of the correlation and variance-covariance matrices. We prefer to use distinct latent variables in each submodel to an approach based on a common latent variable which is scaled by factor loadings in each submodel (eg: Dunson 2000, McCulloch 2008), as the linearity assumption required for the latter is too restrictive for present purposes (Gueorguieva 2001). For the latent variable approach, we do not include response variables as covariates in any of the submodels. The submodels for each stage are presented below, followed by the multivariate distribution of latent variables.

#### 2.1.1 Stimulation phase submodel

For patient *j*, we assume the number of oocytes (eggs) obtained 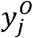 follows a Poisson distribution and model the log of the rate parameter 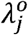 in the usual way:

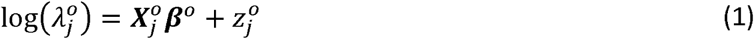

where 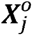 is a row-vector of cycle-level covariates for patient *j, β*^0^ is a corresponding vector of regression parameters and 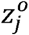 is a patient-specific latent variable that captures overdispersion in the oocyte yield. This submodel is fitted to all patients who start the cycle.

#### 2.1.2 Fertilisation submodel

We model the number of embryos obtained when oocytes are mixed with sperm 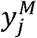 in terms of its rate parameter 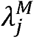, again using a Poisson submodel:

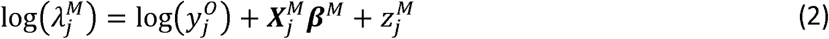

where 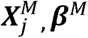 and 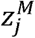 are analogous to the corresponding terms in the stimulation model. We now include an offset term corresponding to the logarithm of the number of oocytes obtained in the linear predictor. This submodel is fitted to all patients who have oocytes mixed with sperm. In some cycles, the number of oocytes mixed with sperm is less than the number obtained, so there is an implicit assumption in the model that any oocytes which were not mixed could not have been successfully fertilized. The assumption is reasonable, since the decision not to mix an oocyte with sperm is almost always based on the fact that the oocyte has been identified as being degenerate.

#### 2.1.3 Embryo quality submodels

We include two measures of embryo quality; cell evenness *(y*^*E*^) and degree of fragmentation *(y*^*F*^). These are ordinal 1 to 4 grading scales measured at the level of individual embryos. We model these using cumulative logit submodels. For embryo *i*(where *i* = 1,2 … *n*_*j*_) nested in patient *j* we have, for *k =* 1,2,3:

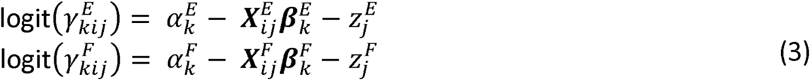

where 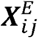 and 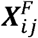 - are row-vectors of covariates 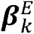 and 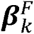 are vectors of regression coefficients which may vary across the levels of *k* (relaxing the proportional odds assumption), and 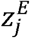 and 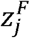 are patient-level random effects (latent variables) which are identified due to the clustering of embryos within patients. 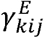 and 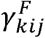 are cumulative probabilities of embryo *i* in patient *j* having a grade of *k* or lower for evenness and fragmentation degree respectively and af and 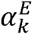 and 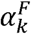 are threshold parameters, corresponding to the log-odds of the embryo having grade *k* or lower. These submodels are fitted to all embryos.

#### 2.1.4 Double embryo transfer submodel

In order to jointly model the binary response DET, denoting the number of embryos transferred, with the other response variables, we use a latent variable representation of a probit regression model (Albert & Chib 1993). Let 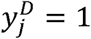 = 1 or 0 if patient *j* does or does not have DET, respectively. We define 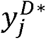 as a latent continuous variable underlying the binary 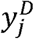 such that:

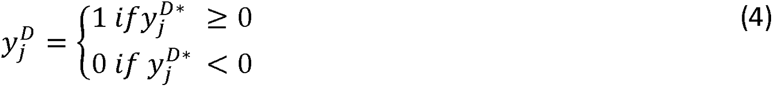

A linear regression submodel for the latent 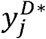 is then used to estimate covariate effects:

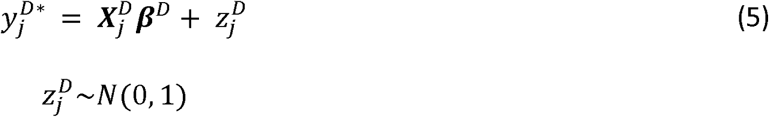

where 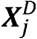 is a row-vector of patient-level covariates and ***β***^D^ is a vector of regression coefficients. Fixing the variance of 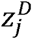 to be 1 is mathematically equivalent to specifying a probit model for the probability that a patient will have DET. We use this error term to link the DET submodel to the others.

#### 2.1.5 Live birth event submodel

As for DET, we use a latent probit representation for 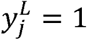 = 1 or 0 corresponding to whether or not LBE obtains, with an underlying latent variable 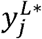 row vector of patient-level covariates 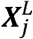 and vector of regression coefficients ***β***^L^. The error term 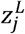 again has a variance of 1, and is used to link the LBE submodel to the others. The DET and LBE submodels are fitted to patients who undergo the transfer procedure.

#### 2.1.6 Covariates in the latent variable approach

An essential feature of the latent variable method is that none of the covariate vectors 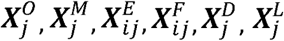 include any of the response variables y_j_.

#### 2.1.7 Latent variable distribution

We specify a multivariate Normal distribution for the latent variables to connect the submodels:

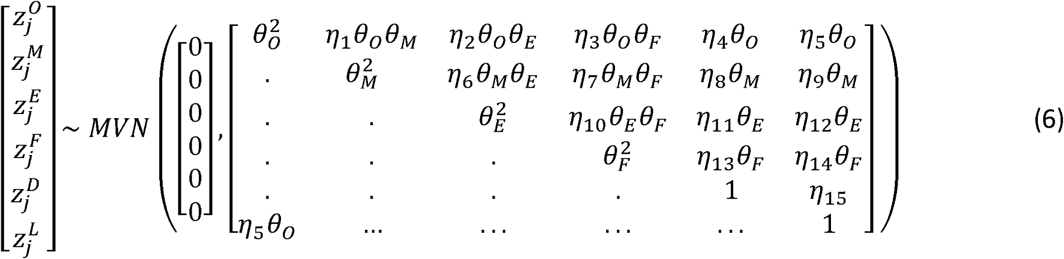

We note that this framework allows us to estimate the relationships between patient and embryo-level responses.

### 2.2 Outcome regression approach

In the outcome regression approach, we fit the submodels presented for the latent variable method as separate regression models, such that we replace the covariance matrix (6) by a diagnonal matrix. By contrast to the latent variable approach, we now include the response variables in the covariate matrices for each downstream submodel. In particular: we include the numbers of oocytes and embryos obtained as covariates in the submodels relating to embryo quality, DET and LBE; we include aggregated measures of embryo quality gradings as covariates in the DET and LBE submodels; and we include DET as a covariate in the LBE submodel. We use the shorthand ‘outcome-covariate’ to refer to instances of response variables appearing as covariates in downstream submodels. While the simplicity of fitting separate regression models makes this an attractive strategy, a weakness of this approach is that it rests on the standard regression assumption that covariates are not *endogenous*, which is to say that they are not correlated with the error term in the submodel. This assumption is unlikely to hold if we include outcome-covariates, due to the likelihood that the different response variables in the submodels share unmeasured predictors.

### 2.3 Combining the latent variable and outcome-regression approaches: an endogenous response model.

A third approach we consider is a combination of the two approaches described above. We represent each response variable using a conditional regression equation including upstream response variables in the covariate matrix, as for the outcome regression approach (2.2). However, we also allow the submodels to be joined through multivariate Normal latent variables as for the correlated latent variable method (2.1). We estimate the variance-covariance matrix of this distribution, together with the regression parameters. This approach allows for the endogeneity of outcome-covariates, since the correlation between response variables is incorporated through the latent variables (Heckman 1978;Terza 1998). Consequently, this approach allows for valid estimation of the effects of upstream upon downstream response variables (Skrondal and Rabe-Hesketh 2004). Identifying endogenous response models can be challenging however (McConnell et al. 2008, Diggle et al. 2007). Standard strategies include fixing parameters in the model (for example, fixing elements of the latent correlation matrix to be zero) and including instrumental variables in some of the submodels (Xie 2000; Steele et al. 2009; Terza 2009). These variables should be strongly correlated with the response variable of the submodel in which they appear, but should not otherwise be associated with downstream responses.

## 3 Application of the methods to routinely collected IVF data.

### 3.1 St Mary’s Data

We utilise the three methods in an application to a routine clinical database from St Mary’s Hospital Department of Reproductive Medicine, Manchester, England. Our aim was to establish whether the endogenous response model would allow us to infer more about the internal structure of the IVF cycle compared to the simpler latent variable and outcome regression methods. The dataset includes 2962 initiated IVF treatments undertaken by 2453 women between 2013 and 2015, including quality data on 12,911 embryos. For present purposes, we ignore the fact that some women underwent multiple cycles, noting that the current models could be extended to a three-level setting (Goldstein 2011). Characteristics of the treatment cycles in the dataset are presented in Table 1. We include age and partner age in all of the submodels. We standardise these by subtracting the mean value and dividing by a standard deviation. The models are flexible enough to allow different covariates to be included in different submodels; we include attempt number in the number of oocytes and DET submodels, pooling 4^th^ and 5^th^ attempts due to small numbers in these categories. In the embryo evenness and fragmentation submodels, we also include an indicator variable denoting whether the egg was fertilized by injecting it with sperm, or by mixing *in vitro.* We suppose that covariate effects are constant across the levels of the ordinal embryo responses (proportional odds), although the methods can accommodate non-proportionality. We fit three models as described in section 2 (correlated latent variable model, outcome regression model, and endogenous response model). Figure 2 shows path diagrams for each of these. Note that we do not distinguish between linear and nonlinear relationships in this diagram.

**Figure 2:**
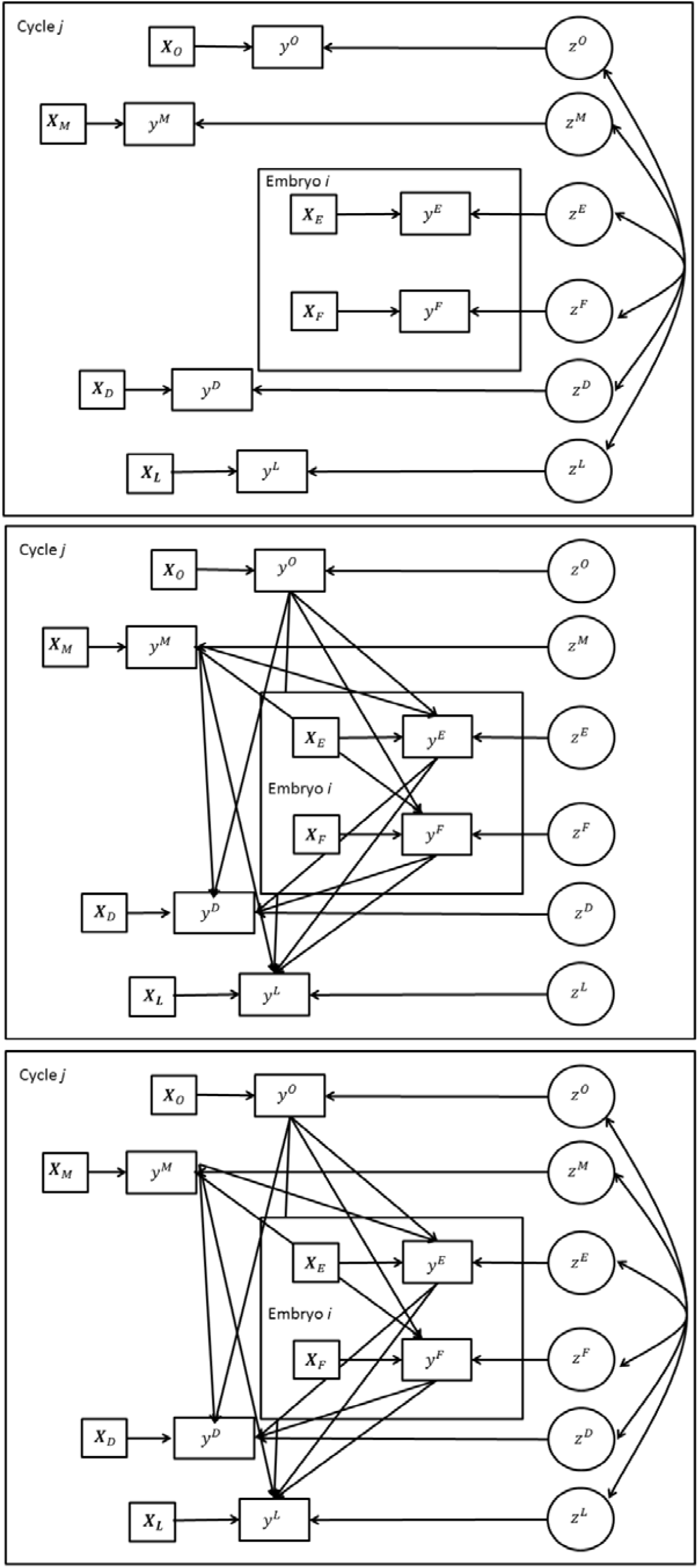
Path diagrams for three models of the IVF cycle: a latent variable model (top), an outcome regression model (middle), and an endogenous response model (bottom).

**Table 1:** Characteristics of the clinical dataset analysed in section 3. Median, interquartile range and range for continuous variables.

| Variable | Summary |
| --- | --- |
| No of cycles started | 2962 |
| No of cycles where eggs mixed with sperm | 2861 |
| No of gradable embryos | 12911 |
| Number of embryo transfer procedures | 2501 |
| Age (years) | 33<br>30 to 36<br>21 to 43 |
| Partner Age (years) | 35<br>32 to 39<br>19 to 72 |
| Attempt Number |  |
| 1 | 2132 (72%) |
| 2 | 659 (22%) |
| 3 | 147 (5%) |
| 4 | 4 (0%) |
| 5 | 20 (0%) |
| Number of eggs obtained per cycle started | 9<br>5 to 13<br>0 to 50 |
| Number of gradable embryos per cycle started | 3<br>1 to 5<br>0 to 19 |
| Number of embryos transferred per transfer procedure | 1049 (42%) |
| 1 | 1452 (58%) |
| 2 |  |
| Live birth event per transfer procedure |  |
| No | 1692 (68%) |
| Yes | 809 (32%) |

### 3.2 Fitting the models

We use the R (R Core Team 2014) implementation of the Bayesian software Stan (Stan Development Team 2014) to fit the models. While the benefits (or drawbacks, depending on one’s perspective) of Bayesian methods have been well rehearsed, our use of this software is primarily driven by pragmatism; the software is flexible and can accommodate complex multilevel models without the need to author custom sampling algorithms. We place weak Normal (0,1000^2^) priors on the regression parameters in the submodels, with the exception of those included in the latent probit submodels (that is, those corresponding to DET, LBE). Given the fact that the latent responses in these submodels have a variance of 1, we place Normal (0, 2^2^) priors on the regression parameters. These can be considered to be weakly informative prior distributions which improve efficiency in fitting the model by restricting the sampler to realistic values for these parameters (Gelman et al. 2014). We place weakly informative Cauchy (0,2.5) priors on the free variance parameters. Finally, we use an LKJ prior distribution for the latent correlation matrix, which is uniform over all possible correlation matrices (Stan Development Team 2017). We consider this appropriate given that estimation of this matrix is of particular interest in our latent variable models. We run the samplers for between two and three thousand iterations in each case, using three chains. We check convergence using the Gelman-Rubin convergence diagnostic (Gelman and Rubin 1992) and using traceplots. In practice, we note that fitting the endogenous response model can take around 12 hours on an Intel Core ¡7-4810MQ 2.8 GHz processor with 16 GB of RAM. Stan code is provided in the online supplement.

### 3.3 Results and interpretation

The models produce a large number of parameter estimates relating to the covariates in each submodel and the relationships between stage-specific responses. In the following, we focus on the information provided by each approach regarding the relationships between response variables, and simply note here that estimates relating to other covariates were generally similar between the models.

#### 3.3.1 Latent variable approach

In the latent variable approach, information regarding the relationships between the variables is obtained through the estimated latent correlation structure (Table 2). We note that estimates derived from this model are generally consistent with current understanding. For example, the model suggests a positive relationship between embryo evenness on the one hand, and probability of LBE on the other (transferring higher quality embryos makes success more likely), although the association between fragmentation and LBE is less clear. The number of oocytes obtained from ovarian stimulation and fertilization rate also appear to be associated with LBE (reflecting advantages of having a larger pool of embryos from which to select). Upstream measures of success are negatively associated with DET, possibly due to the fact that the transfer of multiple embryos is usually employed to compensate for poor prognosis. On the other hand, it is not immediately obvious why fertilization rate is (quite strongly) negatively related to the number of oocytes obtained, and to embryo quality variables.

**Table 2:**
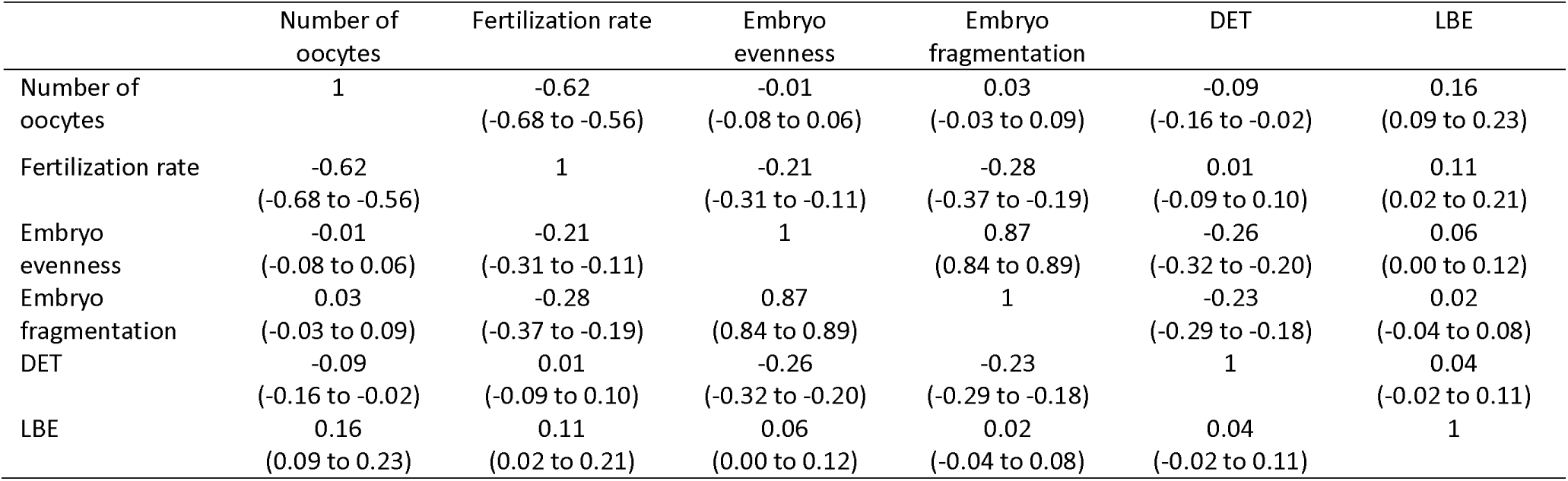
Estimates of association between IVF response variables from the correlated latent variable model. Posterior medians and 95% CIs.

More generally, we might ask whether the latent correlations arising from this approach can reasonably be given a causal interpretation. In relation to this, we note that the estimated correlation coefficients can be adjusted for confounding variables by including these in both of the relevant submodels. However, giving correlation coefficients a causal interpretation may be problematic even if they are appropriately adjusted, since their magnitude is in part determined by the variance of the covariables under consideration, which may vary across populations (Greenland et al. 1991). Moreover, the estimated correlation between any two response variables is not adjusted for other response variables in the model. As a result, it is not possible to distinguish genuine from spurious structural relationships attributable to confounding by other outcome variables. Consequently, the latent variable approach does not appear to yield interpretable estimates of the relationships between response variables. In section 4, we suggest that the latent variable approach may be more useful for the purpose of making multivariate predictions regarding IVF cycle outcomes. We note here that the latent variable approach accommodates both multilevel response data and participants with incomplete response data. Correlations relating to embryo-level responses can be interpreted as measures of association with the patient’s mean values of fragmentation and evenness, while drop out is assumed to be ‘missing at random’ (MAR, Rubin 1976) given the observed responses and covariate data (McCulloch 2008).

#### 3.3.2 Outcome regression approach

The outcome regression approach provides information on relationships between response variables directly by way of estimated regression coefficients (Table 3). Unlike the correlation coefficients from the latent variable model (Table 2), these are adjusted for upstream response variables as well as the other covariates in the submodel. The regression coefficients are also easier to interpret compared to the latent correlations and, moreover, may be given a causal interpretation. In the outcome regression approach, the parameter corresponding to fertilization rate in the LBE submodel is an estimate of the effect of increasing fertilization rates on LBE for a fixed number of oocytes, after blocking effects acting via the intermediate outcomes embryo quality or DET (Westreich and Greenland 2013). The estimate (95% Cl) is 0.14 (0.08 to 0.21), indicating a positive effect. For the estimates in the outcome regression model to be valid however, we must assume that there is no unmeasured confounding (Westreich and Greenland 2013). For example, for our estimates of the effects of embryo evenness and fragmentation in the LBE submodel to be valid, we must assume that there are no unmeasured variables which influence both embryo quality and LBE. This is unrealistic in practice, as there are likely to be deleterious factors which influence both embryo viability and uterine receptivity (Roberts et al. 2010). Even if we believed that this could be adequately accounted for by the inclusion of age as a covariate, residual confounding due to measurement error and model misspecification (for example, including age as a linear term when its relationship with several of the responses may be nonlinear) would ensure that this assumption did not hold (Sterne et al. 2016). In the outcome regression approach, subgroups of participants enter each submodel according to their progress through (and drop out from) the stages of treatment. The model assumes therefore that missing data can be accounted for by the predictor variables in each submodel (and are therefore MAR given these covariates).

**Table 3:** Regression coefficients from outcome regression and endogenous response models for the IVF cycle. Posterior medians and 95% CIs.

| Parameter | Outcome regression model | Endogenous response model |
| --- | --- | --- |
| **Number of oocytes submodel** |  |  |
| Intercept | 2.09 (2.07 to 2.12) | 2.09 (2.07 to 2.12) |
| Age (SDs) | -0.18 (-0.21 to -0.15) | -0.18 (-0.21 to -0.15) |
| Partner Age (SDs) | 0.04 (0.01 to 0.06) | 0.03 (0.01 to 0.06) |
| Attempt number: 1<sup>st</sup> | 0 | 0 |
| 2<sup>nd</sup> | 0.06 (0.01 to 0.12) | 0.07 (0.02 to 0.12) |
| 3<sup>rd</sup> | 0.17 (0.06 to 0.27) | 0.15 (0.06 to 0.24) |
| 4<sup>th</sup> or 5<sup>th</sup> | 0.02 (-0.23 to 0.26) | 0.14 (-0.10 to 0.37) |
| **Fertilization rate submodel** |  |  |
| Intercept | -1.04 (-1.07 to -1.01) | -0.96 (-0.99 to -0.93) |
| Age (SDs) | 0.06 (0.03 to 0.10) | 0.07 (0.04 to 0.10) |
| Partner Age (SDs) | -0.02 (-0.05 to 0.02) | -0.02 (-0.05 to 0.01) |
| **Embryo evenness submodel** |  |  |
| Intercepts (log odds of <=k): k=1 | -4.33 (-4.47 to -4.18) | -4.33 (-4.49 to -4.19) |
| K=2 | -1.37 (-1.47 to -1.28) | -1.37 (-1.47 to -1.27) |
| K=3 | 1.34 (1.24 to 1.43) | 1.35 (1.25 to 1.45) |
| Age (SDs) | 0.02 (-0.04 to 0.09) | 0.02 (-0.09 to 0.13) |
| Partner Age (SDs) | 0.04 (-0.02 to 0.11) | 0.04 (-0.02 to 0.11) |
| Sperm injected into egg | -0.26 (-0.38 to -0.14) | -0.26 (-0.38 to -0.15) |
| Number of oocytes | 0.09 (0.01 to 0.17) | 0.24 (-0.10 to 0.60) |
| Fertilisation rate | -0.16 (-0.23 to -0.09) | -0.45 (-0.66 to -0.21) |
| **Embryo fragmentation submodel** |  |  |
| Intercepts (log odds of <=k): k=1 | -5.07 (-5.26 to -4.88) | -5.07 (-5.25 to -4.88) |
| K=2 | -2.41 (-2.56 to -2.27) | -2.40 (-2.54 to -2.26) |
| K=3 | -0.30 (-0.43 to -0.16) | -0.28 (-0.41 to -0.15) |
| Age (SDs) | -0.12 (-0.21 to -0.03) | -0.22 (-0.37 to -0.07) |
| Partner Age (SDs) | 0.07 (-0.02 to 0.16) | 0.08 (-0.02 to 0.17) |
| Sperm injected into egg | -0.32 (-0.48 to -0.16) | -0.32 (-0.48 to -0.15) |
| Number of oocytes | 0.22 (0.09 to 0.34) | -0.09 (-0.53 to 0.42) |
| Fertilisation rate | -0.30 (-0.41 to -0.20) | -0.57 (-0.90 to -0.20) |
| **Double embryo transfer submodel** |  |  |
| Intercept | 0.13 (0.07 to 0.19) | 0.08 (0.02 to 0.14) |
| Age (SDs) | 0.07 (0.00 to 0.13) | -0.03 (-0.11 to 0.05) |
| Partner Age (SDs) | -0.03 (-0.09 to 0.03) | -0.02 (-0.07 to 0.04) |
| Attempt No: 1<sup>st</sup> | 0 | 0 |
| 2<sup>nd</sup> | 0.25 (0.12 to 0.37) | 0.26 (0.15 to 0.38) |
| 3<sup>rd</sup> | 0.47 (0.23 to 0.70) | 0.53 (0.31 to 0.76) |
| 4<sup>th</sup> or 5<sup>th</sup> | 0.63 (0.08 to 1.22) | 0.68 (0.15 to 1.23) |
| Number of oocytes | -0.06 (-0.13 to 0.02) | -0.25 (-0.48 to 0.01) |
| Fertilization rate | -0.02 (-0.09 to 0.05) | -0.37 (-0.53 to -0.16) |
| Embryo evenness | -0.14 (-0.20 to -0.08) | -0.09 (-0.15 to -0.02) |
| Embryo fragmentation | -0.12 (-0.18 to -0.06) | -0.04 (-0.12 to 0.03) |
| **Live birth event submodel** |  |  |
| Intercept | -0.55 (-0.64 to -0.47) | -0.38 (-0.73 to -0.06) |
| Age (SDs) | -0.06 (-0.12 to -0.00) | -0.12 (-0.20 to -0.04) |
| Partner Age (SDs) | -0.05 (-0.11 to 0.02) | -0.03 (-0.09 to 0.03) |
| Number of oocytes | 0.04 (-0.03 to 0.13) | -0.11 (-0.35 to 0.14) |
| Fertilization rate | 0.14 (0.08 to 0.21) | -0.17 (-0.35 to 0.03) |
| Embryo evenness | 0.07 (0.01 to 0.13) | 0.03 (-0.04 to 0.10) |
| Embryo fragmentation | 0.04 (-0.24 to 0.10) | 0.04 (-0.04 to 0.11) |
| DET | 0.11 (0.00 to 0.21) | -0.15 (-0.63 to 0.44) |

#### 3.3.3 Endogenous response modelling approach

The latent correlation matrix from the fitted model is not obviously interpretable (Web Table 1). Instead, we use the regression coefficients to investigate the relationships between variables (Table 3). As for the outcome-regression model, the regression coefficient corresponding to an outcome-covariate can be interpreted as an estimate of the effect of the outcome on the response variable in the submodel. This estimate applies for fixed values of any upstream (in relation to our outcome-covariate) variables, after blocking indirect effects through intermediate (downstream) response variables. We note that several of the estimates are inconsistent with those obtained from the outcome regression model. For example, the estimate (95% Cl) corresponding to fertilization rate in the LBE submodel changes from 0.14 (0.08 to 0.21) in the outcome regression model to −0.17 (-0.35 to 0.03) in the endogenous response model. This suggests that the positive relationship between fertilization rate and LBE probability observed in the previous models might have been an artefact due to measurement error and unmeasured confounders; in the endogenous response model we conclude that an increased fertilization rate is probably associated with a reduced chance of a successful transfer. This might reflect an increased risk of selecting inferior embryos that are not identified by the grading scales used here. This contrast highlights the possibility of using endogenous response models to explore the extent of unmeasured confounding. In general, the estimates of outcome-covariate effects are less precise in the endogenous response model compared to the outcome regression model. This is analogous to the impact of allowing for, rather than ignoring, clustering of repeated measurements in a mixed model. To check the model, we plotted the observed responses in the data against replicated data drawn from the posterior predictive distribution (Figure 3). For embryo evenness, embryo fragmentation, DET and LBE, we plotted the observed frequency distributions with 95% intervals for the predictive distributions from the model. These checks suggested that the model was compatible with the study data, with the exception of DET, which systematically exceeded the model predictions by a small amount (Figure 3). This is because our prior for the DET intercept was too strong, resulting in underfitting. We would relax this in future applications.

**Figure 3:**
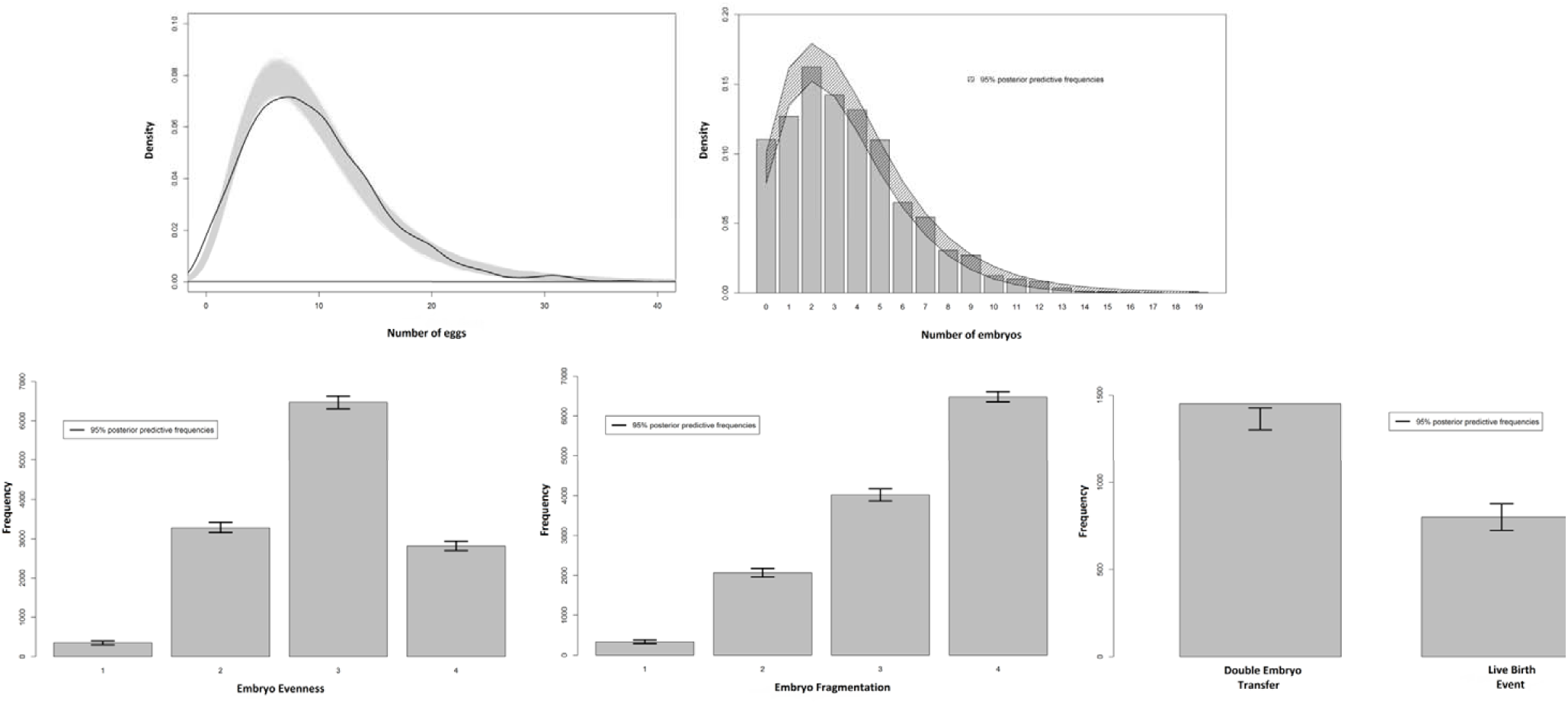
Observed response distribution against replicates drawn from the posterior predictive distribution. ‘Error bars’ on barplots are 95% intervals based or replicated proportions.

The endogenous response model is again a MAR approach, as missing data are assumed to be ignorable given observed response and covariate data. Since the endogenous response model provides interpretable estimates of effects of procedural responses on downstream events (unlike the latent variable model) while allowing the assumption that outcome-covariates are not endogenous to be relaxed (unlike the outcome regression model) we conclude that this approach is superior for the analysis of multistage IVF treatment data.

## 4 Discussion

We have presented and compared several approaches for the analysis of multistage IVF data. All methods offer several advantages over those previously described, including the ability to incorporate mixed outcome types and responses defined at different levels of a multilevel data structure. The approaches are flexible enough to accommodate different combinations of response types and different covariates in the various submodels, according to the particular research question under consideration. The models can be fitted in freely available Bayesian software (Stan Development Team 2017) without the need to write custom sampling algorithms.

The application to routinely collected clinical data highlighted a number of key differences between the approaches. The latent variable method can be used to examine relationships between covariates and stage-specific response variables. However, it is less useful for investigating the relationships between the responses, due to difficulties in interpreting the latent correlation coefficients and the fact that these cannot be adjusted for other response variables in the model. Both the outcome regression approach and the endogenous response model were preferable in this regard. Both provide easily interpretable regression coefficients and allow the causal structure of the sequence of responses to be represented. The validity of the estimates in the outcome regression approach rests upon an assumption of no unmeasured confounding however, which will always be implausible in practice. By contrast, the endogenous response model allows for the valid estimation of outcome-covariate effects by explicitly modelling the correlation between the error term and the response variable (Terza 1998, Skrondal and Rabe-Hesketh 2004). We have assumed a multivariate Gaussian distribution for the latent variables connecting the submodels. This is unverifiable in practice. Accordingly, the model should not be seen as a panacea for confounding. It might be possible to improve robustness in this regard using more flexible latent variable distributions, such as mixtures of Normals (Komarek et al. 2010) or copula-based methods (de Leon and Wu 2011). We are aware that, in discussing the potential of these methods to quantify structural relationships in the IVF cycle, we have skirted the debate about whether or not it is meaningful to speak of causal effects of variables which are not directly modifiable (Greenland 2017, Krieger and Davey Smith 2016). The methods we present could be described using the language of causal mediation, so that instead of speaking of an effect of number of oocytes on downstream variables, for example, we could speak of the effects of predictors being mediated through the number of oocytes. A valid approach to mediation analysis in nonlinear models has been described by Pearl (2011).

Identification of endogenous response models can be challenging. The inclusion of instrumental variables in some of the submodels is a common strategy to assist with identification. In our analysis, ‘attempt number’ acted as an instrumental variable in the number of oocytes and DET submodels and the binary variable ‘method of fertilization’ (either by mixing with sperm *in vitro* or by injecting the sperm directly into the oocytes) acted as an instrumental variable in the embryo quality submodels. We therefore assumed that attempt number affects the number of oocytes obtained and the decision to transfer two rather than one embryos (with previous failed attempts making it more likely both that higher doses of ovarian stimulation drugs will be used and that two rather than one embryo will be transferred) but does not influence the other response variables other than via these intermediaries. We also assumed that the method of fertilizing the oocytes influences the downstream outcomes solely through the quality of the resulting embryos. It is difficult to imagine how the method of fertilization could affect the cycle outcome by any other causal pathway. There could plausibly be unmeasured common causes of our instruments and downstream responses, which would undermine their validity as instruments. We note however that, since we handle endogenity through correlated latent variables in the model, validity of the instruments is not required. We anticipate that identification of endogenous response models is likely to be easier using larger datasets than that considered here, although as noted above fitting the models can be computationally expensive. It remains to identify a suitable reparametrization which may improve the speed of fitting the model, and to investigate the role of Bayesian prior regularization in improving efficiency (Gelman et al. 2014).

Although the latent variable model was not useful for the purpose of investigating relationships between responses, models of this sort can be usefully employed for the purpose of making multivariate predictions (McCulloch 2008). Using the posterior draws from the latent variable model fitted in section 3, we predicted the IVF cycle outcomes for a new cohort of patients with the same characteristics as those in our sample. We adopted a sequential approach whereby we predicted the number of oocytes obtained after stimulation for each patient for each draw from the posterior, before predicting the fertilization rate (and consequently, the number of embryos obtained) in those who were predicted to have any oocytes available. We then predicted the embryo quality for each embryo predicted to arise from the fertilization procedure, and so on. This approach allows us to predict the responses of a cohort of patients (or indeed, of an individual) as they pass through the stages of the IVF cycle, while incorporating the dependency between stage-specific responses. This is not a feature of existing prediction models (eg: Dhillon et al. 2016, Nelson and Lawlor 2011), but may be useful to the clinician whenever there is interest not only in the overall outcome of treatment but also in ensuring that this is achieved in a safe manner. For example, large egg yields following ovarian stimulation are associated with increased risk of ovarian hyperstimulation syndrome (Steward et al. 2014), low birthweight and preterm birth (Sunkara et al. 2015). Consequently, a target of ovarian stimulation is to obtain a yield of oocytes which is neither too low to limit the overall likelihood of success, nor too high to represent a risk to the patient or offspring (La Marca and Sunkara 2014, La Marca et al. 2012). We can use the latent variable approach to predict the probability of a patient obtaining a safe yield of oocytes under a given treatment (eg: fewer than 15) and going on to have a live birth. Without conditioning on any patient or treatment covariates, we calculate this as 23%, with a 95% prediction interval of 21 to 25%. It remains to establish whether there is any advantage offered by including outcome-covariates in the prediction setting.

While all of the models presented here can accommodate embryo-level response variables, relationships between these and other outcomes are estimated using the mean value (Dunson et al. 2003). An undesirable consequence of this is the implicit assumption that the relationship between the evenness and fragmentation of an embryo is the same as the relationship between the evenness of an embryo and the fragmentation of another from the same patient (Gueorguieva 2001). This could be relaxed by using latent representations of the embryo grading submodels and allowing the embryo-level residual terms to be correlated (McCulloch 2008, Gueorguieva and Sanacora 2006). A related concern is the fact the models do not allow embryo-level responses to be included as covariates in the DET and LBE submodels without averaging the values over a patient’s embryos. The estimation of the effects of embryo characteristics on birth outcomes is complicated by the fact that if two embryos are transferred and only one implants, it is not known which of the two was successful. This partial observability problem motivates the use of embryo-uterine models which have been described from both Bayesian (Dukic and Hogan 2002) and Likelihood (Roberts 2007) viewpoints. It remains to incorporate the embryo-uterine approach in the joint modelling approaches described here. We also note that the mean value might not be the best summary measure to use for the purpose of including the embryo gradings as covariates in the DET and LBE submodels, since the best one or two are selected for transfer. An alternative measure capturing the highest available grades might be more appropriate future applications of the methods. Alternatively, we could include the quality of the transferred embryos as additional embryo-level response variables in the model. More generally, we note the fact that we included only a small number of covariates as a limitation of our analysis. We anticipate that differences between the outcome regression and endogenous response approaches will reduce if further covariates are available to control for confounding. This is a topic for future research.

In these examples, we do not differentiate between twin and singleton births. The difference is clinically important, since twin pregnancies represent increased risk to the mother and infants. While we do not make this distinction here, any of the approaches we describe could be extended to accommodate a twin vs singleton submodel, fitted conditional on birth. Our live birth submodel also does not distinguish between failure due to transferred embryos not implanting in the uterine wall and failure due to implanted embryos not being sustained to term (ie: miscarriage). This may be an important distinction for some research questions. Separate submodels for embryo implantation and birth (conditional on implantation) could be included to this end.

All of the approaches presented here assume that any drop out from the cycle is MAR. This might be plausible, since drop out is usually the result of poor response or outright failure at one stage, preventing further progression. These response variables are included in the models. If however, the MAR assumption is deemed not to be appropriate, we could jointly model the drop out process by defining a sequential probit submodel (Albert and Chib 2001) corresponding to transitions through the stages, and allowing this to be correlated with the stage-specific responses (Steele et al. 2009). An alternative strategy would be to employ a selection modelling approach (Heckman 1976, 1978, Diggle and Kenward 1994) where the probability of dropout at a given stage is related to the coincident (possibly unobserved) response variables by way of inclusion as covariates and/or correlated latent variables. Selection models are difficult to implement in Stan, which does not support discrete parameters, thereby precluding explicit modelling of missing egg counts or ordinal gradings. More generally, we might question whether *missing-not-at-random* (MNAR) methods are suitable in the present context. Given that downstream responses are defined conditional on upstream success, this may be construed as an example of the phenomenon known as *truncation-by-death* (Zhang and Rubin 2003, Rubin 2006). McConnell and colleagues (2008) note that MNAR methods implicitly assume an underlying value for missing outcomes, and discuss principal stratification approaches as an alternative. The applicability of principal stratification methods to complex multistage IVF data warrants consideration in future research.

## 5 Recommendations

When analysing multistage IVF data, the appropriate analytic method will depend on the exact research question under consideration. If interest is in estimating the effects of treatment and patient characteristics on outcomes, as well as the structural relationships between the responses at each stage, we recommend the use of the endogenous response model. Identification of the model is likely to require relatively large, detailed datasets, and researchers should be realistic about the scope to answer mechanistic research questions where this resource is not available. Questions of this sort may be tackled using the outcome regression approach, but researchers should be aware that this involves the strong assumption that confounding has been adequately dealt with by measured covariates. In our simple example, we arrived at substantively different conclusions regarding the effects of procedural response variables on downstream outcomes in the outcome regression approach compared to the endogeneity approach, which allows for the correlation between procedural responses and unmeasured predictive factors. We would urge caution when interpreting the endogenous response model however, since inevitable misspecification of the latent variable distribution means that residual confounding will not be eliminated. Researchers should still attempt to reduce confounding through the inclusion of known variables as far as possible. Estimates corresponding to other (exogenous) covariates were similar between models. Where interest lies in making predictions about the patient journey through the stages of the IVF cycle, the relatively simple latent variable approach offers a framework to do this while taking the dependency between the stages into consideration. These approaches assume that drop out is MAR. We are unable to make a definitive recommendation regarding the most appropriate approach to modelling drop out at present, other than to state that MNAR methods assume that there is an underlying value for each missing response. This may not be appropriate where responses are strictly undefined. Finally, given the complexity of IVF, we note that any meritorious analysis will require substantial input from clinician and clinical scientist collaborators.

